# A spinal substrate for modular control of natural behavior

**DOI:** 10.64898/2026.03.12.711392

**Authors:** Fabricio Nicola, Lily Li, Tiernon Riesenmy, Randall Pursley, Robert Brian Roome, Jeffrey E. Markowitz, Ariel J. Levine

## Abstract

Natural behavior unfolds as coordinated sequences of body movements. This organization suggests that behavior may be built from discrete motor patterns, yet how such arrangements are implemented by neural circuits remains unknown. Here, we combined kinematic analysis, muscle recordings, genetically identified cell types, and closed-loop optogenetic perturbations to examine the organizational logic of natural gap-crossing jumps in mice. Jumping was characterized by a series of precisely defined phases and their associated modular motor patterns. The core phases, propulsion and flight, exhibited distinct signatures of neural control, including unique bursts of coordinated hindlimb muscle activity, differential tuning strategies for jump distance, and active requirements for spinal neural drive. Mapping activity across lumbar interneuron populations and functionally screening candidate cell types for their ability to drive coordinated movement revealed that a population of dorsal excitatory dILB6 neurons can autonomously evoke coordinated multi-joint hindlimb flexion characteristic of the jumping flight phase, across behavioral contexts. These findings provide a specific cellular substrate for the long-standing concept of spinal modular motor control: a flexible, preconfigured motor template that the mammalian CNS can recruit and modulate to meet the demands of natural behavior.

**Highlights:**

- Jumping behavior is organized into discrete phases with stereotyped motor patterns
- Each phase contains a distinct signature of body-wide, joint, and muscle coordination
- dILB6 spinal interneurons autonomously evoke a triple-flexion hindlimb motor module
- dILB6 neurons can shape active jumping behavior

## Introduction

Animal behavior evolves as organized sequences of coordinated body-wide configurations across space and time ^1–3^. Even simple actions emerge from the integration of many biomechanical degrees of freedom, yet they proceed as coherent multi-joint movements. Quantitative analyses of muscle activity, kinematics, and posture have formalized this organization by identifying recurring patterns of motor output across different scales, such as co-activation of muscle groups (“synergies”) and recurring body-wide action patterns (“syllables”)^4–15^. The remarkable reproducibility of these motor patterns across behaviors and animals supports the influential view that natural behavior may be composed from a foundational set of distinct motor elements, or modules ^16–21^. In this study, we use the term “module” to refer to a coherent pattern in which multiple neurons, muscles, joints, or body regions are recruited together as a unit. The coordinated action of these potentially independent components could reflect a common control input, such as a shared neural signal or (bio)mechanical force. However, how modularly organized behavior could be implemented by the mammalian nervous system remains a fundamental problem in motor control. In particular, it is unclear whether stereotyped motor behavior reflects an underlying modular organization of neural control, and, if so, where and how such modules are instantiated within motor circuits.

To investigate this question in mice, we sought a behavior from their natural repertoire that might be decomposed into separable phases and provide insight into motor control across diverse contexts. We focused on gap-crossing jumps, part of a family of jumping behaviors that includes the vertical hunting pounce of a red fox, the acrobatic aerial parkour of squirrels, and the nimble leaps of a mountain goat ^22–26^. Like many goal-directed behaviors, jumping requires specific movements to occur in a particular order over time to achieve a successful outcome, such that propulsion must precede flight, and landing necessarily follows both. This non-rhythmic temporally ordered structure could facilitate the study of how multiple movements are organized within a behavior.

Support for separable movement modules for jumping comes from both evolutionary and developmental processes. Across vertebrates, features of the jump were incrementally added to the behavior: basal frogs jump by extending both hindlimbs and then land in a belly flop ^27^. Evolutionarily newer amphibians perform the same hindlimb extension, followed by hindlimb flexion and controlled landings using the forelimbs to absorb the impact ^27^. Reptiles, birds, and mammals added the preparatory “counter-movement” that enhances take-off ^28–31^, and mammals progressively elaborated these patterns, adding optional elements such as forelimb swing during propulsion or complex landing strategies ^32,33^. In mice, jumping emerges naturally during postnatal development, around eleven days after birth, when pups can support their body weight and exhibit spontaneous lunging, followed a few days later by vigorous high-velocity limb extensions (“popcorn jumps”) ^34–36^, but it takes days to weeks of subsequent maturation and practice to achieve the precisely controlled, phased jumps that are characteristic of adult animals ^36^. The progressive appearance of distinct features of jumping supports the idea that gap-crossing jumps could be composed of separable neural control units, making this behavior a promising test case for examining modular organization in goal-directed mammalian movement. Moreover, aspects of jumping behavior are preserved in some spinalized animals (in which a surgical lesion separates the brain from the spinal cord and body) ^4,37–41^, suggesting that the spinal cord may serve as a node for coordinating such motor patterns.

Here, we asked (1) whether natural jumping behavior in mice is indeed organized into discrete motor modules, (2) whether specific spinal interneuron populations can generate such modules, and (3) whether these spinal circuits shape ongoing jump execution by combining kinematic analysis with closed-loop optogenetic manipulations of specific spinal cord cell types. We first analyzed the body-wide kinematic structure of gap-crossing jumps and identified distinct behavioral phases. A focused analysis of propulsion and flight revealed that these phases are likely mediated by distinct neural control mechanisms: multi-joint extension for propulsion and multi-joint flexion for flight. We then mapped jumping-associated spinal neurons, optogenetically screened candidate cell types for their ability to drive jumping modules, and found that a specific dorsal excitatory population, dILB6 interneurons, can autonomously evoke context-independent, multi-joint hindlimb flexion when mice are at rest, in propulsion, or mid-air during flight. Our results reveal that natural jumping comprises distinct motor modules and identify a specific population of spinal interneurons that can evoke a triple-joint module and shape ongoing natural behavior.

## Results

### Natural jumping behavior exhibits modular organization

Rodents can cross gaps several times their body length in a behavior typically described as unfolding over a preparatory period with active visual distance estimation, followed by a stereotyped sequence of propulsion, flight, and landing ^23,33,42,43^. Although this phase-based description is widely adopted in behavioral and neurophysiological studies, it remains unclear whether these divisions reflect the intrinsic organization of the behavior or are imposed by the experimenter’s perception of movement categories. To address this, we performed video recording, markerless tracking of body landmarks, and analysis of the temporal evolution of body movement without imposing predefined phase boundaries, allowing the intrinsic structure of behavior to emerge directly from the data.

Jumping was highly successful and reproducible across animals and trials, consistent with prior work ^42,43^. 36 of 39 animals performed consistent horizontal jumps between the platforms, while three animals exhibited escape-like movements and were excluded from further analysis (Supplementary Fig. 1A). In the remaining 36 animals, we observed the previously described visually-defined phases (Fig. 1A). Mice first adopted a preparatory crouched posture near the edge of the takeoff platform, followed by coordinated body-wide extension during propulsion leading to takeoff, hindlimb flexion during flight, and forelimb-assisted landing.

**Figure 1.**
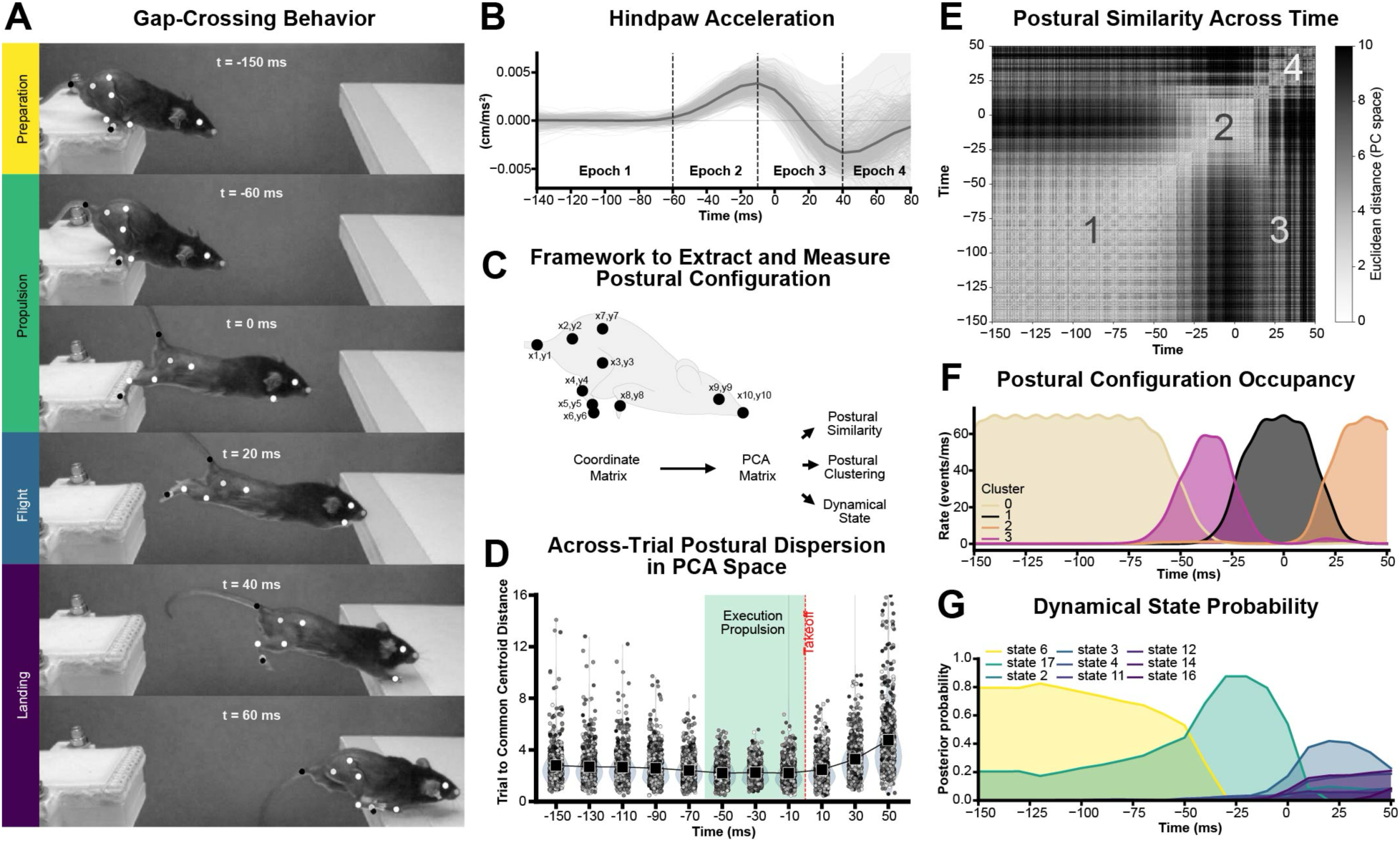
Whole-body kinematics reveal discrete postural and dynamical states during natural jumping. (A) Gap-crossing jump behavior. Representative frames from high-speed video illustrating the stereotyped sequence of postural configurations during a gap-crossing jump. Key body landmarks used for kinematic tracking are shown as white points. The behavior unfolds through a sequence of phases: preparation, propulsion, flight, and landing. (B) Hindpaw acceleration across the jump sequence. Thin grey lines show individual trials. The thick line indicates the mean acceleration, and the shaded region indicates ±1 s.d. across trials (n = 36 subjects, 695 trials). Vertical dashed lines indicate boundaries between behavioral epochs identified from kinematic dynamics. (C) Framework to extract and quantify postural configuration. Body posture was represented as a coordinate matrix of tracked anatomical landmarks. Postural configurations were projected into a low-dimensional space using principal component analysis (PCA), enabling quantification of postural similarity, clustering of recurrent configurations, and identification of dynamical behavioral states. (D) Across-trial postural dispersion in PCA space. Trial-to-centroid distance of posture trajectories aligned to takeoff. Postural variability is low during the propulsion phase, indicating stereotyped limb configurations during movement execution. (E) Postural similarity across time. Time-by-time matrix of Euclidean distances in PCA space showing similarity between postures across the jump sequence. Numbers mark regions of high postural similarity corresponding to epochs where body posture is stable and geometrically constrained, separated by rapid transitions. (F) Postural configuration occupancy. Temporal occupancy of postural clusters across the jump, showing discrete configurations that dominate specific time windows. (G) Dynamical state probability. Posterior probability of dynamical states inferred using an autoregressive hidden Markov model (AR-HMM). Distinct states are sequentially engaged during the jump, reflecting structured transitions in the behavioral dynamics.

To begin assessing whether jump execution exhibits intrinsic biomechanical structure independent of visual phase labels, we tracked hindpaw acceleration as a representative point on the mouse hindlimb during jumping. Across animals and trials, we found that the acceleration profile exhibited clear phases rather than a gradual shift (Fig. 1B). Paw acceleration remained near baseline at the start of the movement, rose quickly before takeoff, dropped suddenly when the limb left the platform, decreased during flight, and increased again at landing. These quick changes split the apparently continuous behavior into four distinct phases, providing direct quantitative support for the intuitive, visually identified preparation, propulsion, flight, and landing.

Jumping is a body-wide movement, so to reveal the comprehensive patterns of motor organization and assess their potential continuous or modular organization, we next analyzed a low-dimensional representation of postural configuration across ten body landmarks (Fig. 1C). For each frame, the x–y coordinates of tracked landmarks were assembled into a coordinate matrix and subjected to principal component analysis (PCA), generating a postural state space that captured the dominant geometric variations of the body. This framework allowed us to (1) quantify postural similarity and variability across time and trials, (2) characterize whether body configurations are organized as separable modes, or clusters, and (3) reveal whether the dynamic movements of jumping are organized as separable action “states”.

Variability in movement execution is thought to reflect underlying control strategies, prompting us to examine how this important motor parameter was organized over time. We measured the distance of individual postures to a common centroid in PCA space across trials (Fig. 1D) and found that postural variability (dispersion among individual postures) was lowest during propulsion (-50 to -10 ms), indicating strong convergence onto common postural configurations during this phase. In contrast, dispersion was higher during preparation (-150 to –70 ms) and during landing (with a higher median), peaking at 50 ms (β = 2.52, p < 0.001). These data indicated that motor variability itself varies over the course of the behavior, in a pattern aligned with the preparation, propulsion-flight, and landing phases of the jumping, even though this analysis did not impose any predefined temporal segmentation. In addition, we found that propulsion represented the most reproducible and geometrically constrained pattern of jumping behavior.

We next examined the patterns of body configuration and how they evolved over time to determine whether they changed gradually or in discrete steps, consistent with continuous or modular control organization, respectively. We computed pairwise Euclidean distances between postures across all time points within the behavioral execution window (Fig. 1E) and observed a striking block-like pattern. If body configuration changed smoothly and evenly, we would expect gradual changes in similarity across time – a blurred diagonal line. Instead, we found that postures that were close in time formed coherent regions of high similarity, separated by sharp changes at specific moments in the behavior. Thus, postural configurations remain stable for finite periods before rapidly shifting to new regions of postural state space. The matrix organization also revealed off-diagonal regions in which temporally distant time points exhibited notable similarity. This most likely reflected a recurring flexed hindlimb posture adopted during both preparation and flight. Overall, these kinematic data show that the mouse body adopts a temporal series of stable, repeatable whole-body configurations separated by rapid transitions, a signature of modular behavior organization.

To characterize the particular body configurations that make up gap crossing behavior, we clustered the postural data and analyzed the presence of each body configuration cluster over time (Fig. 1F). Leiden clustering at a relatively low resolution produced four robust clusters that each corresponded to a single phase of jumping behavior (Supplementary 1D). In temporal order: cluster 0 showed a flexed hindlimb posture with a shortened trunk and was characteristic of preparation, cluster 3 showed an elongated body posture with forelimbs extended but hindlimbs flexed and was characteristic of the first half of propulsion, cluster 1 showed a fully extended posture for the forelimbs, trunk, and hindlimbs and was characteristic of propulsion and the onset of flight, and cluster 2 showed extended forelimbs with a shortened trunk and flexed hindlimbs and was characteristic of flight and the preparation for landing. Overall, we observed an orderly sequence of modular, whole-body postural configurations.

While the instantaneous body postures revealed the “spatial” patterns of jumping behavior, they do not capture the dynamic, “temporal” patterns of movement. To determine whether jump execution occurs as a series of distinct movement states, we used an autoregressive hidden Markov model (AR-HMM) on the postural data. We observed a small number of movement states, with their probabilities changing abruptly over time (Fig. 1G). At the start of the movement, a dominant preparatory state gave way to a propulsive state that peaked just before takeoff, followed by a transition, after which a group of states related to flight and landing appeared. We found that even varying AR-HMM parameters, state probabilities consistently concentrated around the same constrained body configurations identified by the similarity matrix and clustering analyses, indicating that these postures represent stable landmarks in the behavioral repertoire (Supplementary 1F). In general, the transitions between states further suggest that the behavior switches between distinct modes.

Together, this kinematic analysis revealed that jumping has a modular architecture composed of a sequence of stable, geometrically constrained whole-body configurations separated by rapid movement transitions. This organizational pattern is consistent with previous studies suggesting that complex actions may be composed of smaller coordinated elements ^5,7,16,44^.

### Propulsion and flight modules reflect different neural control signatures

We hypothesized that the modular organization of jumping behavior reflects an underlying modular structure of neural circuits, with distinct circuits controlling distinct movements. In particular, the “core” modules of gap crossing - propulsion and flight, could be mediated by the prototypical hindlimb muscle synergies: multi-joint extension and multi-joint flexion. To assess whether propulsion and flight are likely to be under separable neural control logic, we performed a focused analysis of hindlimb muscle activity during these phases, examining whether patterns differ across task variations and whether each phase requires active neural drive.

We first quantified joint angles as an indirect measure of summed muscle forces at the hip, knee, and ankle (Fig. 2A-B). Across animals and trials, joint trajectories exhibited highly stereotyped temporal profiles. During preparation and early propulsion, the knee and ankle flexed, a hallmark of the “counter-movement” that precedes powerful jumps in mammals and birds ^28,31,45–47^. All three hindlimb joints then underwent rapid, dramatic extension, reaching their peak angles just after takeoff, when the platform’s ground reaction forces no longer act on them. Immediately after, all three hindlimb joints flexed for the duration of flight, followed by partial re-extension in preparation for landing.

**Figure 2.**
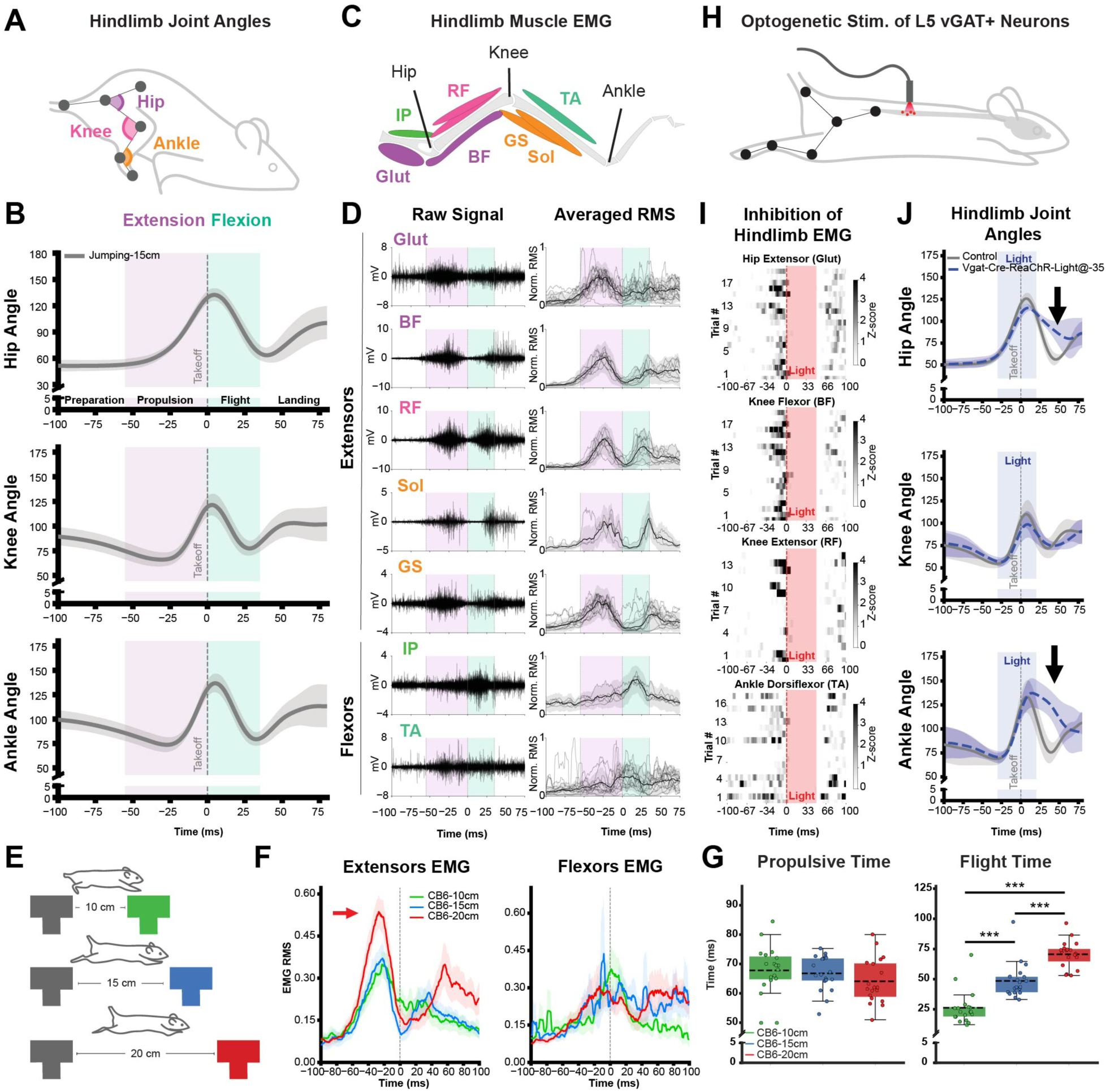
Propulsion and flight phases exhibit distinct hindlimb coordination and neural control signatures. (A) Schematic illustrating hindlimb joint angles (hip, knee, ankle) extracted from tracked anatomical landmarks during voluntary jumping. (B) Mean hindlimb joint-angle trajectories (± s.d., n = 36) during a 15-cm jump. Shaded regions denote behavioral epochs (preparation, propulsion, flight, landing). Purple and green shaded regions indicate propulsion and flight, respectively. Vertical dashed line indicates take-off. (C) Schematic of recorded hindlimb electromyography. Extensor muscles include gluteus (Glut, n=15), biceps femoris (BF, n=8), rectus femoris (RF, n=9), soleus (Sol, n=3), and gastrocnemius (GS, n=12); flexor muscles include iliopsoas (IP, n=5) and tibialis anterior (TA, n=13). (D) Raw electromyography (EMG) traces (left) and trial-averaged root-mean-square (RMS) EMG signals (right) for hindlimb muscles, aligned to take-off. Shaded regions indicate extension and flexion epochs as in (B). Traces averaged within each mouse are shown in gray; bold traces indicate the mean across mice. (E) Experimental schematic illustrating different jump distances (10, 15, and 20 cm) used to modulate task demand. (F) Mean flexor and extensor EMG RMS activity across jump distances. Shaded regions indicate ± s.e.m. Arrow highlights the distance-dependent modulation of muscle activity. Compared with the 10 cm, peak extensor activity at 20 cm increased (β = 0.269, p < 0.001, mixed-effects model; n = 18–19 animals per group). (G) Quantification of propulsion and flight durations across gap distances. Flight time differed significantly across gap conditions (one-way ANOVA, F_2,54_ = 53.15, p < 0.001). Tukey’s post hoc test indicated significant differences between all gap distances (all p < 0.001). (H) Experimental schematic illustrating optogenetic stimulation of L5 spinal vGAT-positive neurons during jump execution. (I) Trial-by-trial heat maps showing normalized EMG activity (z-scored RMS) for selected hindlimb muscles, aligned to optogenetic stimulation. Vertical red bars indicate the timing of light delivery. (J) Mean hindlimb joint-angle trajectories (± s.d.) during control trials and trials with optogenetic stimulation of L5 vGAT-positive neurons delivered 35 ms before take-off. Blue shading indicates the light pulse. Arrows highlight phase-specific deviations in joint kinematics. Hip and ankle joint angles differed between conditions at specific timepoints during flight (t-tests at each timepoint, p < 0.001), whereas knee angles did not differ significantly (n = 6 controls, n = 5 stimulation).

To reveal the particular muscle activity patterns that mediate these joint movements, we recorded electromyography activity from primary hindlimb flexor and extensor muscles spanning the hip, knee, and ankle joints (Fig. 2C). Across jumps, individual muscles exhibited reproducible activation profiles, with consistent timing and waveform shape evident in both raw EMG traces and trial and mouse-averaged RMS signals (Fig. 2D). Muscle recruitment was strongly phase-specific, similar to prior work in other animals. ^44,48^ First, the extensor muscles showed coordinated low-level activation, indicating that the flexed hindlimb posture observed during preparation reflected an eccentric contraction, in which the muscle was active during lengthening. Eccentric contractions are thought to increase the force created by subsequent concentric contractions of the same muscles by activating the 1a proprioceptive reflex loop, by placing the muscle in an optimal place on the force-length curve, and by storing elastic energy in tendons. ^49–51^ Next, the extensor muscles exhibited a large burst of activity preceding propulsion, likely providing the primary takeoff force. After this phase, the hip flexor (iliopsoas) and ankle flexor (tibialis anterior) muscles displayed a distinct and highly stereotyped pattern of co-activation preceding flight. Last, the extensor muscles contracted again in preparation for landing. Bi-articular muscles of the hindlimb cross two joints and typically act as flexors at one joint and as extensors at the other, with the dominant action depending on the joint angle ^52–54^. We found that the bi-articular muscles rectus femoris, biceps femoris, and gastrocnemius were each activated together with the extensor burst preceding propulsion, likely reflecting maximal recruitment of hindlimb muscles to amplify extensor force generation.

Overall, hindlimb muscle activity closely mirrored joint kinematics and revealed a clear dual-module organization: an extensor burst for propulsion, and a flexor burst for flight. The simplicity, synchrony, and consistency of these coordinated extension and flexion activation patterns are consistent with the previously proposed “muscle synergy” hypothesis, in which a common pre-motor neuron drive (low-dimensional neural control) coordinates multiple (high-dimensional) muscle activity ^21,55–57^.

Rather than serving as rigid building blocks, muscle synergies are thought to be modulated depending on context ^5^. To test whether the modules in propulsion and flight phases could be flexibly tuned to task demands, we systematically varied gap distances and examined how kinematics and muscle patterns were modulated across small, medium, and large jumps (Fig. 2F-G; Supplementary video 1). Mixed-effects modeling revealed that increasing jump distance significantly increased the magnitude of the extensor bursts during the propulsion phase (Fig. 2F; mixed-effects model, p < 0.001), whereas flexor bursts during flight remained unchanged. Increasing distance was also associated with longer flight duration, as mice briefly remained suspended during the longest jumps, while propulsion duration did not change significantly. These results indicate that increasing task demand does not alter the fundamental coordination pattern of the jump. Instead, longer jumps are achieved by scaling the magnitude of extensor activation during propulsion while preserving the overall modular structure of the behavior.

Not all aspects of movement necessarily arise from active neural drive; passive biomechanical forces such as inertia, gravity, and limb dynamics can also shape body motion. ^58–60^ To determine whether the stereotyped kinematic and muscular coordination patterns described above depend on active spinal neural drive, we transiently inhibited lumbar spinal circuits at L5 at specific phases of the jump execution (Fig. 2H-J). This manipulation profoundly disrupted hindlimb muscle activity, abolishing EMG signals across multiple muscles spanning all three hindlimb joints during the light pulse. Heatmap representations of the EMG activity revealed a complete suppression of activation in gluteus (Glut), biceps femoris (BF), rectus femoris (RF), and tibialis anterior (TA) under optical stimulation (Fig. 2I). When spinal inhibition was delivered during propulsion, animals failed to complete the jump, indicating that this phase required sustained active neural drive. When the stimulus was delivered during the flight phase, it revealed differential joint vulnerability: while the knee joint-angle excursions remained intact, those of the hip and ankle were profoundly altered (Fig. 2J). These findings indicated that jumping is governed by a hybrid neural-mechanical control, in which active neural drive interacts with passive biomechanical forces, while spinal activity was necessary to drive both the propulsion and flight phases.

Together, we found that propulsion and flight each displayed a different coordinated burst of muscle activity, were tuned differently as the jumping distance increased, and each required active neural drive to maintain their normal joint-angle patterns. These findings suggest that the propulsion and flight modules of gap jumping behavior are mediated by phase-specific, separable neural mechanisms.

### Jumping behavior recruits spatially and molecularly defined lumbar spinal neurons

The extension and flexion synergies can be generated by the spinal circuits ^4,16^, suggesting that the neural substrates underlying these jumping movements may lie within the spinal cord. Previous activity-dependent mapping studies have shown that spinal interneurons are recruited in a behavior-specific way, with distinct patterns along the rostro-caudal and dorso-ventral laminar axes ^61,62^.

To identify jumping-associated neurons, we mapped expression of the immediate early response gene cFOS throughout the lumbar spinal cord. Jumping animals showed significantly higher cFOS density in specific laminae than in controls, with enrichment in the deep dorsal and intermediate regions of the cord. Differences relative to naïve were observed at L2 (laminae V and VIII) and L4 (laminae VI-VIII), whereas relative to the platform control were found at L2 (laminae VII-VIII) and L6 (laminae II-IV, VI-VII, X) (Fig. 3B; one-way ANOVA with Tukey’s test, p < 0.05). In contrast, spinal cords from control animals that were handled and placed on the jumping platform but not allowed to jump showed lower overall cFOS expression with a more diffuse distribution, while naive home cage animals exhibited minimal activation across all segments and laminae.

**Figure 3.**
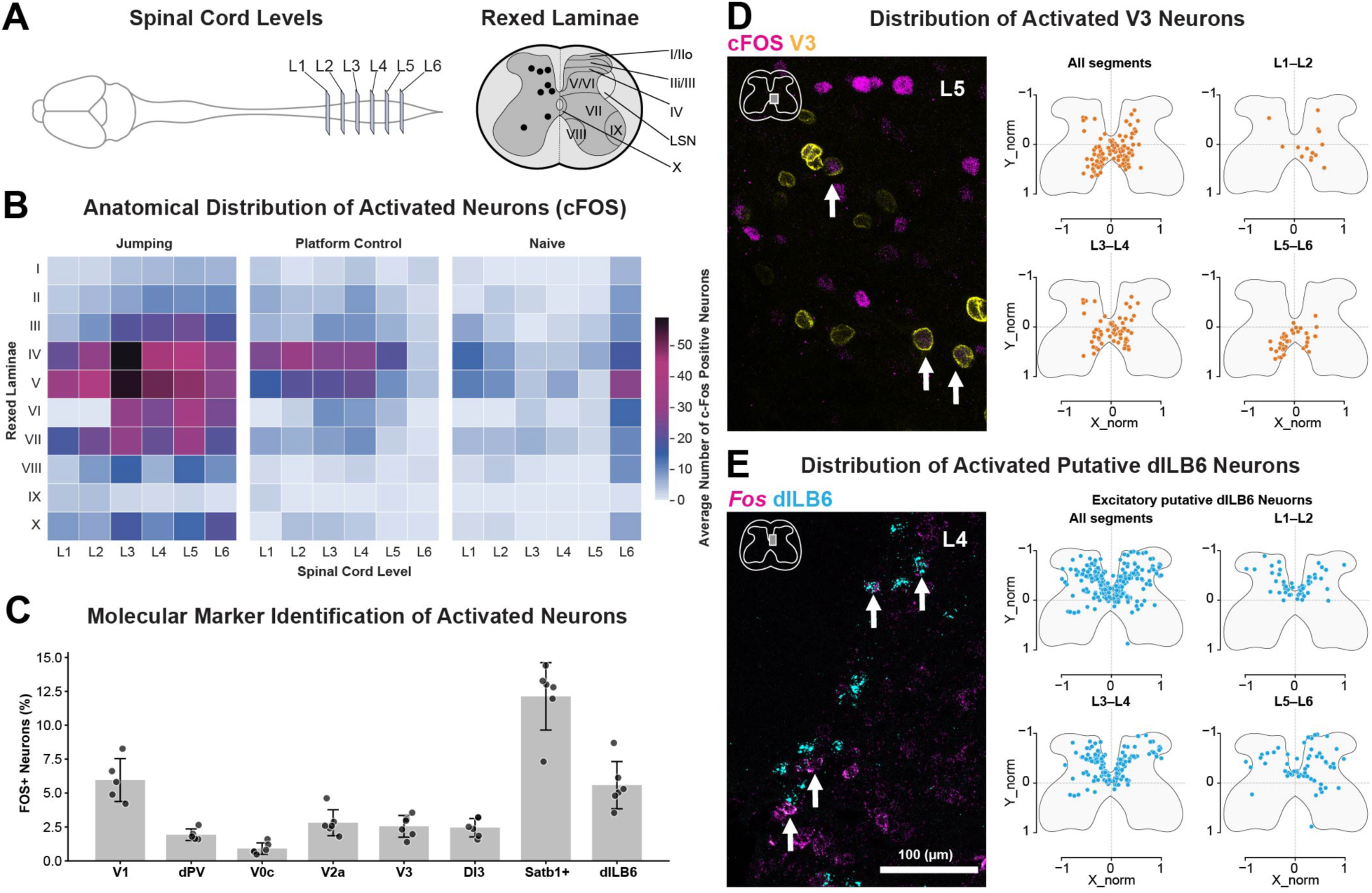
Spatial and molecular identity of lumbar spinal interneurons activated during jump execution. (A) Schematic of lumbar spinal cord segments analyzed (L1–L6) and Rexed laminae used to map neuronal activation. (B) Anatomical distribution of activated neurons across spinal segments and laminae following jump behavior, measured by cFos expression. Heat maps showing the average number of cFos-positive neurons across Rexed laminae and lumbar levels for animals performing gap-crossing jumps, platform control animals, and naïve animals. Significant differences were detected between Jumping and Naïve animals at L2 (laminae V, VIII) and L4 (laminae VI-VIII), and between Jumping and PControl at L2 (laminae VII-VIII) and L6 (laminae II-IV, VI-VII, X). One-way ANOVA followed by Tukey’s test, p < 0.05. n = 6 Jumping, 6 PControl, 2 Naïve. (C) Molecular identity of activated spinal neurons. Quantification of cFos/*Fos*-positive neurons co-expressing molecular markers corresponding to major spinal interneuron classes (V1, dPV, V0c, V2a, V3, and dI3) and excitatory populations, including Satb1+ and putative dILB6 neurons. Data represent the percentage of marker-positive neurons that were cFos/*Fos*-positive. (D) Distribution of activated V3 neurons. Left, representative confocal image showing cFos expression co-localized with V3 neurons in an L5 spinal cord section; arrows indicate double-labeled neurons. Right, normalized spatial distributions of activated V3 neurons across lumbar segments (n=6 animals). Each point represents the position of a labeled neuron within the spinal cord cross-section. (E) Distribution of activated putative dILB6 neurons. Left, representative confocal image showing *Fos* expression co-localized with dILB6 neurons in an L4 spinal cord section; arrows indicate double-labeled neurons. Right, normalized spatial distributions of activated excitatory putative dILB6 neurons across lumbar segments (n=6 animals). Each point represents the position of a labeled neuron within the spinal cord cross-section.

To determine whether the neurons activated during jumping corresponded to specific populations of molecularly defined interneurons, we combined cFOS protein or *Fos* mRNA labeling with markers for defined spinal neuronal populations, focusing on cell types within the peak region of cFOS expression including the inhibitory V1 (*En1*^Cre::INTACT^) and PV^deep^ (*Lbx1*^Cre^;*Pvalb*^FLPo;^*ROSA26*^ds:tomato^) populations and the excitatory V0c (Chat+), V2a (Chx10+), V3 (Sim1^Cre::INTACT^), dI3 (Isl1+), and dILB6 populations (subset of Satb1+ and *Cdh23*+/Slc32a1-) ^63^. Each of these showed expression of cFOS/*Fos* after jumping behavior, with a notable fraction of jumping-associated neurons being dILB6 neurons (Fig. 3C). Although Satb1 labels a large fraction of activated neurons, this marker spans several heterogeneous interneuron populations ^64^, making it difficult to target as a single functional class. We therefore focused on the more molecularly defined dILB6 subset.

To further examine the spatial organization of activated interneurons, we analyzed the distribution of cFOS-positive neurons within a pair of excitatory populations. Activated V3 interneurons displayed a stereotypical spatial pattern, located mainly in ventromedial regions of the lumbar spinal cord and scattered cells in the medial deep dorsal horn, consistent with the known distribution of the overall V3 population (Fig. 3D).^65^ Activated dILB6 *Cdh23*+/*Slc32a1*-interneurons were found mainly in the medial deep dorsal horn of the lumbar cord, in particular in a necklace of neurons surrounding the dorsal funiculus (Fig. 3E). Together, these experiments presented several candidate neuronal populations for mediating aspects of jumping behavior.

### dILB6 spinal interneurons reshape jump behavior via a modular pattern

To identify spinal interneuron populations capable of engaging movement patterns relevant to jumping, we performed optogenetic stimulation of multiple genetically defined interneuron classes while animals were at rest (Fig. 4A-B). We targeted populations identified through activity mapping during jumping. Across the populations tested, focal optogenetic activation produced a range of postural and kinematic responses, including localized muscle contraction, whole-body postural changes, hindlimb movements, and no movement (Fig. 4B and Supplementary Video 2). The V1 (*En1*^Cre::ReaChR^), PV^deep^ (*Lbx1*^Cre^;*Pvalb*^FLPo^*;ROSA26*^ds:ReaChR^), V0c (*Pitx2*^Cre::ReaChR^), and V2a (Vsx2^Cre;^AAV:ReaChR), populations elicited limited or variable effects that did not resemble hindlimb movements observed during jumping, while optogenetic stimulation of V3 interneurons (*Sim1*^Cre::ReaChR^), consistently produced robust hindlimb extension involving the hip, knee, and ankle, leading animals in a crouched posture to transiently extend their hindlimbs (Fig. 4C-D). This triple-joint extension pattern resembled the extension module expressed during the propulsion phase of jumping and is consistent with prior work demonstrating a role for V3 in amplifying activity in hindlimb extensor muscles ^66^. However, none of these spinal populations evoked hindlimb flexion.

**Figure 4.**
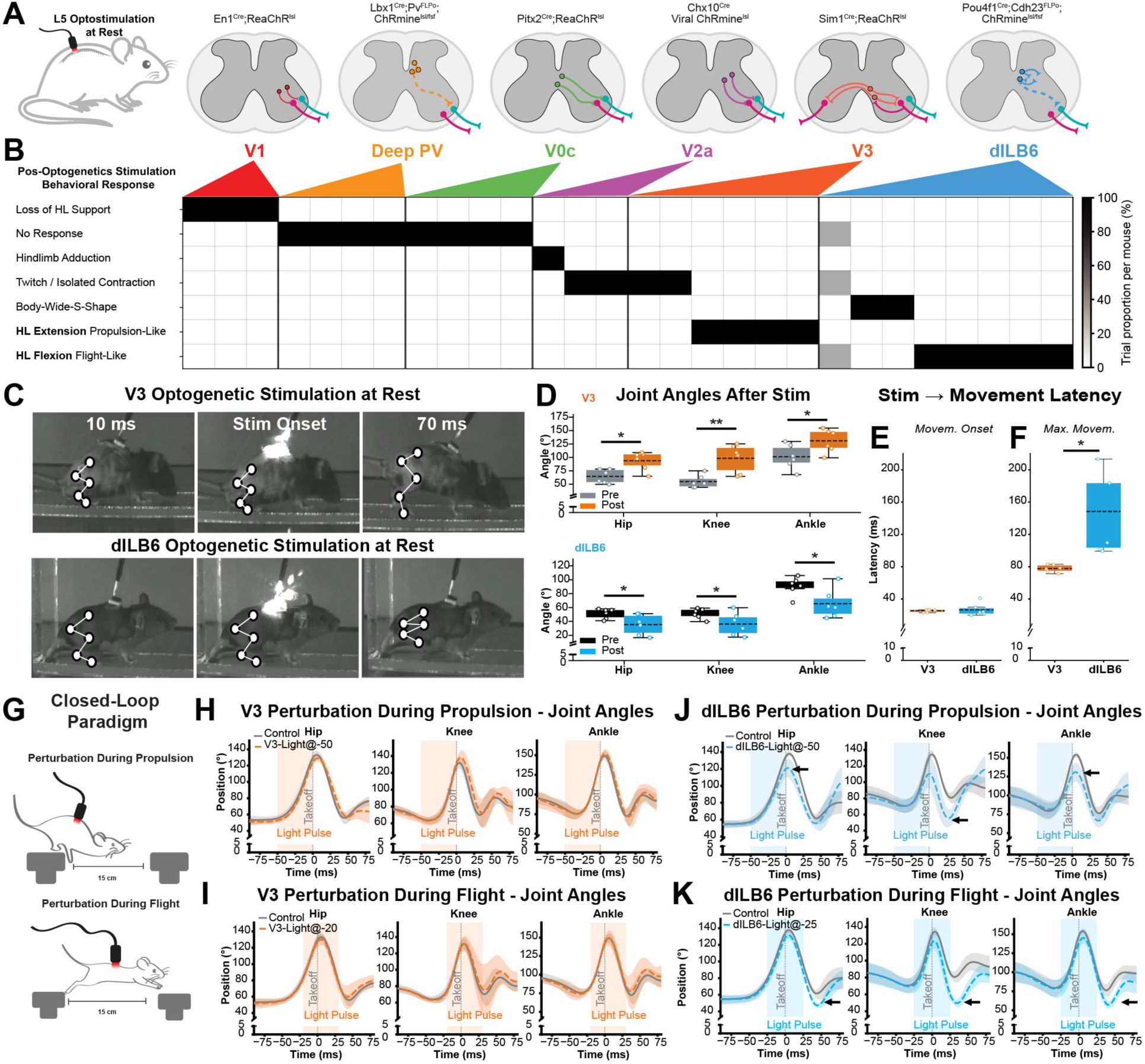
dILB6 spinal interneurons drive a coordinated hindlimb flexion module that reshapes jump execution. (A) Experimental schematic illustrating optogenetic stimulation of genetically defined spinal interneuron populations at the L5 level. Targeted populations include V1 (En1^Cre^), deep parvalbumin-positive interneurons (Lbx1^Cre^;Pvalb^Flp^), V0c (Pitx2^Cre^), V2a (Chx10^Cre^), V3 (Sim1^Cre^), and dILB6 (Pou4f1^Cre^;Cdh23^Flp^), each expressing ReaChR or ChRmine as indicated. (B) Behavioral responses evoked by optogenetic stimulation at rest for each interneuron population. The heat map indicates the proportion of trials per mouse exhibiting distinct motor outputs (columns). (C) Example video frames showing body posture before, at onset, and following optogenetic stimulation at rest for V3 (top) and dILB6 (bottom) interneuron populations. White markers indicate tracked hindlimb landmarks. (D) Quantification of hindlimb joint angles before (Pre) and after (Post) optogenetic stimulation at rest for V3 and dILB6 populations. Box plots summarize animal-level mean joint angles for each condition; individual points represent animals. Statistical comparisons were performed within each group using paired two-tailed t-tests (*p < 0.05, **p < 0.01, n = 6). (E) Latency from stimulation onset to movement initiation response for V3 and dILB6 interneuron stimulation. (F) Latency from stimulation onset to maximal postural response for V3 and dILB6 interneuron stimulation. Each point represents one animal; bars indicate mean ± s.d. (n = 6). Groups were compared using a two-tailed Welch’s t-test (t = 3.41, *p < 0.05; Hedges’ g = 1.82). (G) Closed-loop perturbation paradigm used to deliver optogenetic stimulation during specific phases of the jump. (H–I) Effects of V3 activation during propulsion and flight on hindlimb joint kinematics. Mean joint angles over time are shown with shaded s.d. No significant differences were observed between conditions (n=5). (J) Effects of dILB6 activation during propulsion. Mean joint angles ± s.d. are shown. At 0ms (propulsion), significant differences were observed in hip angle (Welch’s t-test: t = 2.68, p < 0.05, n = 5) and ankle angle (t = 3.52, p < 0.05, n = 5). At 20ms (flight), a significant difference was observed in knee angle (t = 4.07, p < 0.05, n = 5). (K) Effects of dILB6 activation during flight. Mean joint angles ± s.d. are shown. At 20ms (flight), significant differences were observed in hip angle (Welch’s t-test: t = 2.61, p < 0.05, n = 5) and knee angle (t = 3.79, p < 0.05, n = 5). At 40ms (flight), a significant difference was observed in ankle angle (t = 3.95, p < 0.01, n = 5).

We hypothesized the dILB6 neurons could provide a spinal cellular basis for coordinated multi-joint flexion on the basis of (1) their anatomical location in medial laminae V/VI, previously associated with “flexor reflex afferent” pathways ^67–69^, and (2) their expression of *Satb1*, a marker of last-order premotor “motor synergy encoder” neurons that can drive activation of multiple motor groups ^70^, and (3) the significant co-expression of Cdh23 and Fos in excitatory neurons following jumping behavior. dILB6 neurons are late-born, dorsal excitatory spinal neurons that we recently described in embryonic development^63^ and that correspond to Excit-23 and Excit-25 in a harmonized atlas of postnatal spinal cord cell types ^64^. To gain genetic access to these cells, we used a combination of *Pou4f1*^Cre^ and a new *Cdh23*^FLPo^ line we generated for this purpose, together with double-stop AAV injected into the lumbar L5 spinal cords of double-transgenic mice to achieve locally restricted genetic control. Viral reporter expression was mostly restricted to medial laminae IV-VI, with scattered superficial expression in medial laminae I and IV, consistent with the predicted location of dILB6 neurons in spatial transcriptomic data (Supplementary Figure 2B). This approach also revealed the anatomy of these neurons, including a prominent short-range intraspinal projection in the lateral deep dorsal horn, through the dorsal horn bundle of Cajal (Supplementary Figure 2F-G).

To test the function of dILB6 neurons, we optogenetically stimulated them and observed consistent flexion of the hip, knee, and ankle joints, leading the animals to lift their legs. In animals in which a unilateral response occurred, the hindlimb flexion resembled a protective withdrawal movement, raising the possibility that the scattered superficial dILB6 neurons were driving a pain response. However, stimulated animals did not exhibit any signs of distress, such as paw licking, guarding, orbital tightening, or avoidance, responses that are typically observed together with reflex movements upon optogenetic activation of spinal cell types that transmit pain-related signals ^71–74^. Thus, dILB6 neurons were sufficient to evoke coordinated hindlimb flexion in resting animals, a movement that was similar to the motor output of withdrawal reflexes, and to the flexion module expressed during the flight phase of jumping behavior (Fig. 4C-D).

Quantification of the joint angles at the onset and after the stimulation confirmed that the V3 activation significantly increased the extension at the hip, knee, and ankle joints, whereas dILB6 activation significantly increased flexion at the hip, knee, and ankle joints (Fig. 4C-D). In both cases, movement onset followed light stimulation with short latency, indicating a rapid influence on hindlimb coordination. Interestingly, the latency to maximal joint excursion following light differed between the two populations, with dILB6 activation producing a more prolonged hindlimb movement compared to V3 activation (Fig. 4 E-F). Thus, these results identify V3 and dILB6 interneurons as candidate populations capable of engaging in coordinated hindlimb movement patterns that resemble the modules expressed during the propulsion and flight phases of jumping behavior.

We next asked whether perturbing these candidate interneurons during ongoing jumping could influence the execution of the behavior. To test this, we used closed-loop optogenetic stimulation to deliver a brief light pulse (50 ms) to the L5 lumbar spinal cord during defined epochs of gap-jumping, aligned either with the propulsion phase or with the subsequent flight phase (Fig. 4G). Despite their strong ability to evoke hindlimb extension at rest, optogenetic activation of V3 interneurons during jumping did not alter hindlimb joint kinematics during either propulsion or flight (Fig. 4H–I). In contrast, activation of dILB6 interneurons selectively altered hindlimb kinematics. Stimulation during the propulsion phase reduced hip and ankle extension (Fig. 4J). When stimulation was delivered during the flight phase, dILB6 activation produced a pronounced increase in hindlimb flexion, resulting in decreased hip, knee, and ankle angles (Fig. 4K).

Together, we characterized spinal neurons activated by jumping behavior, screened six distinct populations for their potential to elicit coordinated movements associated with jumping, and found that stimulating dILB6 neurons drove coordinated multi-joint hindlimb flexion when animals were at rest, performing the propulsive phase of jumping, or mid-air during flight. Accordingly, these excitatory local spinal interneurons could represent the cellular basis for the flexion muscle synergy and a core module of jumping behavior.

## Discussion

To understand how the mammalian nervous system implements natural behavior, we focused on a highly conserved, ethologically relevant task: gap crossing jumps in mice. We found that jumping is organized into discrete execution phases, each implemented by a distinct set of motor modules and each displaying a different signature of underlying neural control. Next, we identified a genetically defined class of spinal interneurons capable of generating and imposing the coordinated multi-joint hindlimb flexion characteristic of the flight phase, across distinct contexts, including during fast, natural jumping. Our work supports a model in which complex behavior is organized into a sequence of execution phases, each composed of a distinct combination of motor elements, and establishes a specific spinal interneuron population as the neural substrate for a common motor module.

The idea that “lower” motor centers contain neural circuits capable of generating coordinated movements is long-standing. It emerged from early observations that stimulation of the mammalian nervous system, particularly the spinal cord and spinal nerves, could evoke coordinated multi-joint reflexes resembling natural actions ^4,75–78^. In mice, we now know that distinct neural populations in the brainstem are specialized for actions such as reaching versus food handling ^79,80^, or for head orientation ^81^, body turning ^82^, and halting locomotion ^83^, demonstrating that coordinated motor patterns can be embedded within subcortical circuits. Even spinalized animals can perform coordinated hindlimb movements, such as withdrawal reflexes, locomotion, jumping, and scratching, particularly under conditions of strong sensory drive or neuromodulation ^4,37–40,77,84,85^. It is known from cellular ablation experiments that specific spinal cord interneurons are required for proper motor execution ^86–94^. However, whether genetically defined spinal interneuron populations are sufficient to impose coherent multi-joint modules during natural behavior was unresolved.

To identify spinal neurons that may be capable of driving multi-joint motor outputs, we screened and compared the outputs elicited by multiple genetically identifiable classes that were activated during jumping, including V1 inhibitory interneurons (known to be required for normal extension movement during locomotion ^89^); PV deep inhibitory dorsal horn neurons (known to be required for normal extension to flexion transition during locomotion ^94^); V0c cholinergic interneurons (known by enhance recruitment of fast firing motoneurons ^95^); V2a excitatory interneurons (known to be required left-right limb coordination ^86^), and V3 excitatory neurons (known as hindlimb extensor modulators ^66^). We found that excitation of most of these classes elicited nonspecific movements or subtly influenced hindlimb coordination, consistent with broad or overlapping roles in spinal circuit modulation. The ability of V3 neurons to rapidly elicit hindlimb extension and prompt standing in resting mice is intriguing, but the action of V3 neurons was context-dependent and more likely reflects their role as gain amplifiers of extensor muscle activity.

In contrast, activation of dILB6 neurons consistently evoked coordinated multi-joint hindlimb flexion across behavioral contexts, including when animals were at rest, engaged in hindlimb extension, or even mid-air. This context-independent recruitment of a structured coordination pattern, similar to the effects produced by “command-like” neurons, suggests that dILB6 neurons provide access to a preconfigured motor module embedded within spinal circuits. Work in Drosophila has identified genetically defined neurons in the ventral nerve cord, including specific populations such as MGT, wPN1, and 23B ^96–98^, that can trigger stereotyped behavioral actions. These neurons are often described as “command-like” because they initiate complex behavioral programs composed of multiple phases and movements. Our findings suggest that genetically defined neurons in mammalian spinal circuits may operate at a different level of this hierarchy, providing access not to entire behaviors or multiple phases but to individual motor modules that implement coordinated multi-joint movements. These observations imply that nervous systems across phyla contain specialized neurons capable of accessing distinct levels of motor organization, ranging from complete behavioral programs to the modular coordination patterns that compose them.

Even in the context of a simple muscle synergy or motor module, no cell type can function alone. dILB6 interneurons are centered in medial lamina IV/V within the internal basal nucleus of Cajal, a region characterized by extensive integrative connectivity ^99,100^. Neurons in this region receive convergent input from descending pathways, primary sensory afferents, and local spinal circuits ^100–103^. Their axons enter the dorsal horn bundle and extend across multiple segments, forming abundant collateral projections onto other spinal interneurons ^99,102–104^. This broad central and peripheral nervous system convergence onto the dILB6 region and the pattern of dILB6 axonal projections suggest that dILB6 neurons are anatomically positioned to integrate distributed signals, cooperate with ongoing feedback, and impose coordinated multi-joint output across segments. Such connectivity would enable dILB6 neurons to drive complex, context-dependent forms of multi-joint flexion. Indeed, the original conceptions of modular motor control, set forth by researchers such as J. Hughlings Jackson, Charles Sherrington, Nikolai Bernstein, and Emilio Bizzi proposed hierarchical and integrative systems in which lower-level modules are recruited, weighted, fractionated, and modulated by higher levels of behavioral control ^5,21,55,75^. In this view, supraspinal centers select and modulate (parameterize or tune) pre-existing motor modules rather than directly drive individual muscle activity. The tunability of the coordinated motor patterns we observed, including graded modulation across execution phases, is consistent with such a hierarchical control model. These findings could suggest that the spinal cord, and dILB6 neurons in particular, provide a structured pattern solution for motor execution, while descending inputs shape the timing, gain, and contextual deployment of these modules.

By providing a quantitative description of modularity in a natural behavior, this work highlights a key advantage of such an organizational logic: the potential for separable neural “regimes” that are each specialized to control the most important tasks during a defined phase of the behavior ^5^. We propose that gap-crossing jumps in mice involve a cortically controlled preparatory phase ^43,105^, a constrained and powerful propulsive phase driven by brainstem and spinal pre-extensor circuits ^40,41,106^, a flight phase coordinated by dILB6 spinal neurons, and a variable, sensory-guided landing phase ^33,107,108^. By identifying a spinal interneuron class capable of eliciting the multi-joint movement of the flight phase, this work helps to move modular motor control from a descriptive concept to a circuit-resolved substrate in behaving mammals. Most broadly, our results raise the possibility that the spinal cord contains a range of specialized elements for diverse movements that could be flexibly recruited across contexts. Understanding how such modular substrates are organized and engaged in natural behavior could open new avenues for restoring coordinated movement after neurological injury, where reactivation or reweighting of preserved motor elements could support functional recovery ^109^.

## Acknowledgements

We gratefully acknowledge Peter Romanienko at the Genome Editing Shared Resource at Rutgers University who generated the Cdh23:FLPo mice, and the generosity of colleagues who shared mouse lines with us: Jay Bikoff (En1^Cre^)^110^, Victoria Abraira (*Lbx1*^Cre^;*Pvalb*^FLPo^) ^94^, Ying Zhang (*Sim1*^Cre^) ^111^, Tudor Badea (*Pou4f1*^Cre^) ^112^.

## Author Contributions

This project was conceptualized and designed by FN and AJL. The experimental apparatus was built by FN and RP. Experiments were carried out by FN and LL and were analyzed by FN, TR, RR, JEM, and AJL. Figures were made by FN. The manuscript was written by FN and AJL and edited by all authors. Funding was secured, and research was supervised by JEM and AJL.

## Supplemental information

Document S1. Figures S1-S2

Video S1. Gap-Crossing Jumps Across Increasing Distances

Video S2. Motor Responses Evoked by Optogenetic Activation Across Spinal Interneuron Populations

## Methods

### Animals

All experimental procedures were conducted in accordance with the guidelines of the National Institutes of Health and were approved by the National Institute of Neurological Disorders and Stroke Animal Care and Use Committee (ACUC) under protocol number 1384. Mice were housed in a temperature- and humidity-controlled environment on a 12-hour light–dark cycle with food and water available ad libitum. Experiments were performed in adult mice aged ∼2-4 months old at the time of testing. Both male and female animals were included. Control animals consisted of wild-type C57BL/6 or CB6F1/J mice. To target genetically defined spinal interneuron populations, the following Cre and intersectional driver lines were used: V1 interneurons (En1^Cre^), deep parvalbumin-positive interneurons (Lbx1^Cre^;Pvalb^Flp^), V0c interneurons (Pitx2^Cre^), V2a interneurons (Chx10^Cre^), V3 interneurons (Sim1^Cre^), and dILB6 interneurons (Pou4f1^Cre^;Cdh23^Flp^). Sample sizes for each experimental group are reported in the corresponding figure legends.

### Generation of the Cdh23:FLPo mouse line

Cdh23:FLPo targeting was done by CrispR-assisted knock-in to insert a FLPo-WPRE-SV40pA cassette into exon 3 of the Cdh23 gene, in frame with, and 4 amino acids downstream of, the initiating methionine. The donor construct comprised 685 bp 5’ homology arm and 540 bp 3’ homology arm flanking the CrispR guide sequence (TGGTACTTGCTATGCTTG). The line was generated in a C57BL/6 background and maintained on a mixed 50:50 C57B/6; BALB/c background to avoid vestibular phenotypes due to a spontaneous mutation in the Cdh23 gene in C57BL/6 mice.

### Gap-crossing behavioral apparatus

The jumping apparatus consisted of two elevated platforms, each 20 cm above the ground, coated with a silicone rubber sheet to provide traction. Animals initiated jumps from a small “take-off” platform rectangle (10 cm long; 6 cm base width) oriented toward the landing platform. The landing platform measured 30 cm in length and 20 cm in width and could be positioned at variable distances from the take-off to control jump difficulty. The apparatus was illuminated from above using diffuse overhead lighting to ensure consistent visibility of the animal during behavioral recording. All surfaces surrounding the apparatus were covered with green paper to create a uniform background and facilitate visual separation between the animal and the environment during video-based tracking.

### Behavior training

Mice were trained to perform voluntary gap-crossing jumps between the take-off and landing platforms. Animals were motivated to cross by moving from the smaller take-off platform to the larger landing platform. Mice were first habituated to the apparatus and trained to step across small gaps separating the platforms. Over successive trials, the gap distance was progressively increased until animals reliably performed jumping movements. Once animals could reliably jump a 15 cm gap, they underwent structured training consisting of 20 trials per session, with two sessions per day for two consecutive days prior to testing. During training sessions, gap distances were varied from 5 to 20 cm in 5-cm increments. Each trial began with the animal placed on the take-off platform facing the landing platform. Trials ended when the animal successfully landed on the landing platform. Trials in which animals failed to initiate a jump or jumped in a direction other than toward the landing platform were excluded from analysis.

### Statistics

Statistical analyses were performed using custom Python scripts. The specific statistical tests used for each experiment are indicated in the corresponding figure legends. Sample sizes represent individual animals unless otherwise stated. Detailed descriptions of behavioral analysis, kinematic processing, EMG processing, and statistical models will be provided in a subsequent version of this manuscript.

