## Supplementary material for "A spinal substrate for modular control of natural behavior": Document S1

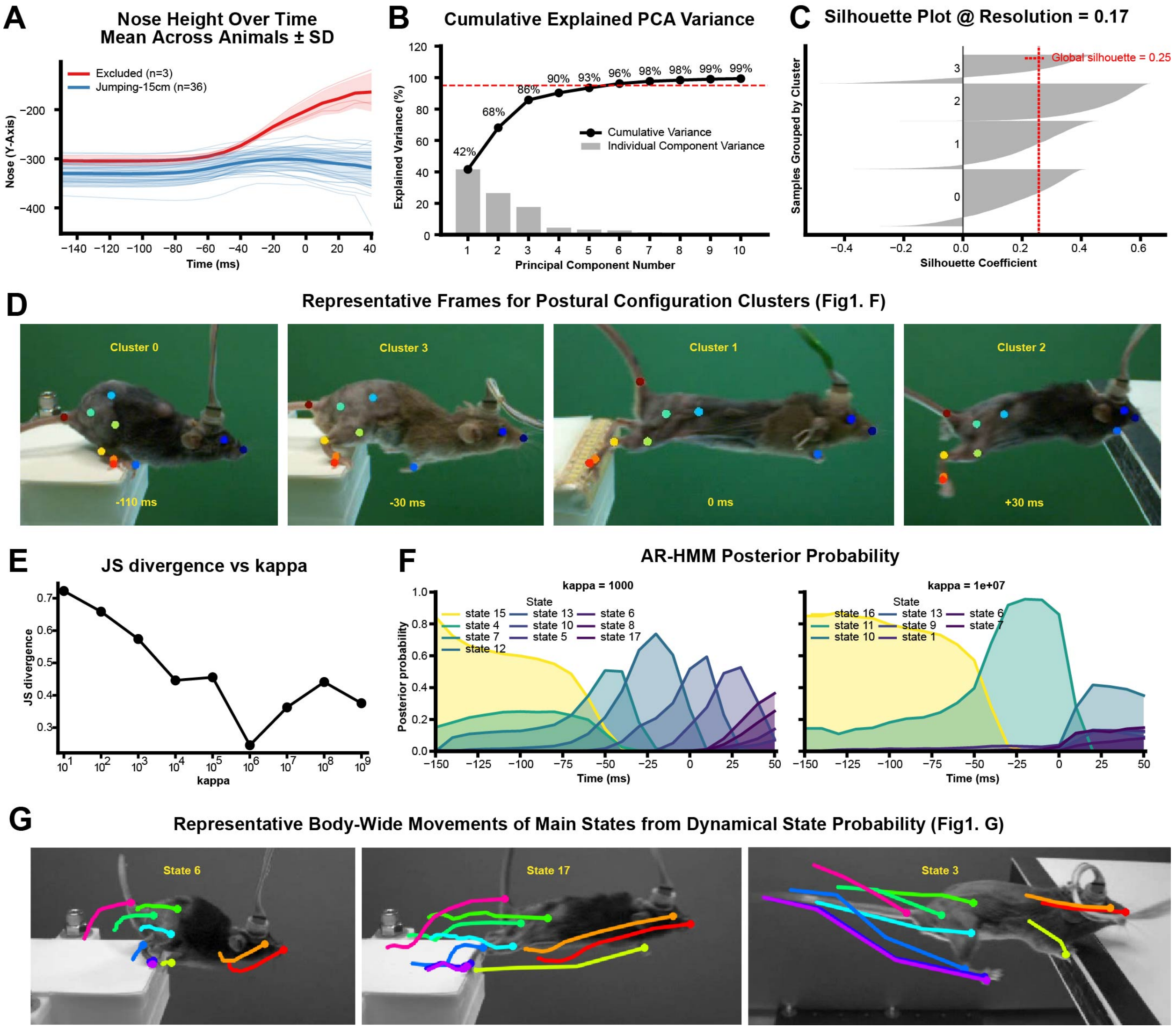


**Figure S1. Identification and Validation of Postural Configuration Clusters and Dynamical States**
(A) Nose height across time (mean ± s.d.). Thin lines show individual trials. (B) Cumulative variance explained by principal components of the posture matrix used for dimensionality reduction. (C) Silhouette analysis evaluating clustering quality for postural configurations. (D) Representative frames corresponding to posture clusters shown in Fig. 1F. (E) Jensen–Shannon divergence across AR-HMM hyperparameter values (κ), used to assess model stability. (F) Posterior probability of AR-HMM states under different κ values, demonstrating robustness of inferred dynamical states. (G) Representative of main body-wide movements corresponding to dynamical states depicted in Fig. 1G.

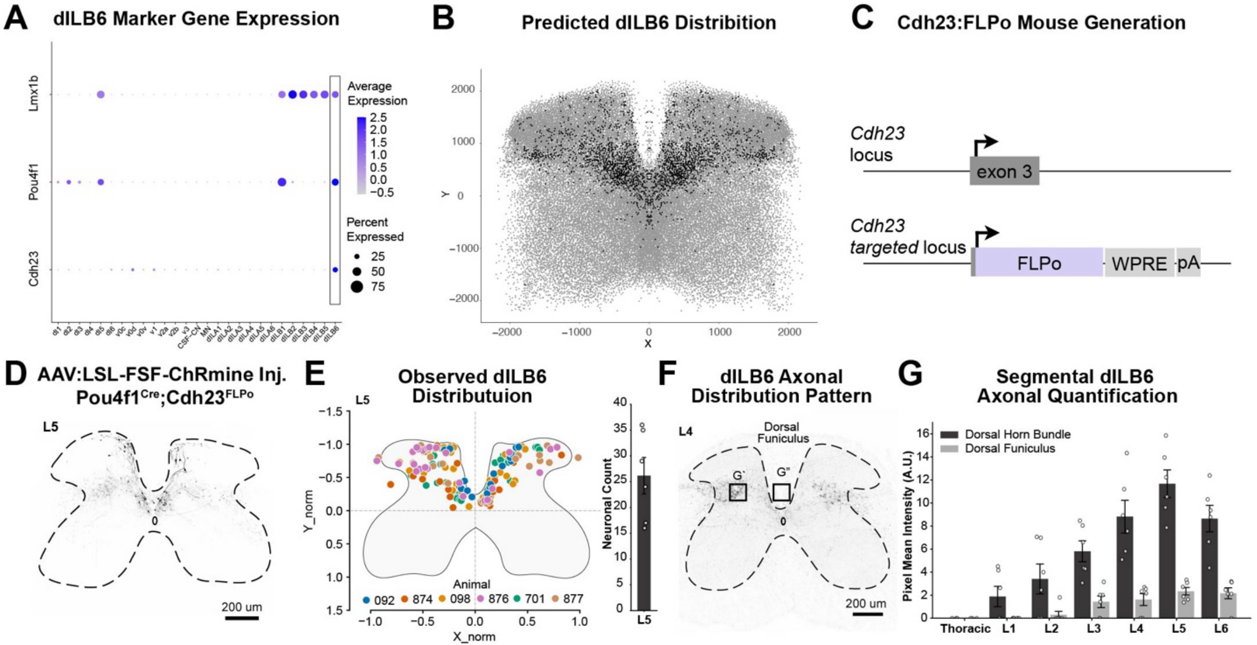


**Figure S2. Genetic Targeting and Anatomical Characterization of dILB6 Spinal Interneurons**

(A) Expression of candidate marker genes used to identify dILB6 neurons from transcriptomic datasets. Dot size indicates the percentage of cells expressing each gene, and color indicates the average expression level. (B) Predicted spatial distribution of dILB6 marker gene expression detected by Xenium spatial transcriptomics in lumbar spinal cord tissue. Each point represents a cell with detected transcripts corresponding to the dILB6 marker gene set, revealing the spatial organization of candidate dILB6 neurons within the spinal cord cross-section. (C) Generation of the Cdh23-FLPo mouse line used for selective targeting of dILB6 neurons. The targeting strategy inserts FLPo recombinase into the Cdh23 locus. (D) Example spinal cord section showing labeling of dILB6 neurons following AAV-LSL-FSF-ChRmine injection in Pou4f1Cre;Cdh23FLPo mice. (E) Observed spatial distribution of labeled dILB6 neurons in lumbar spinal cord sections. Each point represents a labeled neuron; colors indicate individual animals. (F) Distribution pattern of dILB6 axonal projections in lumbar spinal cord sections. Boxes indicate regions used for axonal quantification in G. G`: Dorsal horn bundle; G": Dorsal Funiculus. (G) Quantification of segmental axonal density of dILB6 projections across thoracic and lumbar spinal segments. Bars represent mean ± s.e.m.
